# Role of geography and climatic oscillations in governing into-India dispersal of freshwater snails of the family: Viviparidae

**DOI:** 10.1101/476820

**Authors:** Maitreya Sil, N. A. Aravind, K. Praveen Karanth

## Abstract

The Indian subcontinent has experienced numerous paleogeological and paleoclimatic events during the Cenozoic which shaped the biotic assembly over time in the subcontinent. The role of these events in governing the biotic exchange between Southeast Asia and Indian subregion is underexplored. We aimed to uncover the effects the collision of the Indian and Asian plate, marine transgression in the Bengal basin as well as the paleoclimatic changes in the subcontinent and adjoining regions, on the dispersal of freshwater snail family Viviparidae from Southeast Asia (SEA) to Indian subregion. Extensive sampling was carried out throughout the Indian subcontinent to capture the current diversity of the targeted lineages. Three mitochondrial and two nuclear markers were sequenced from these samples and combined with published sequences to reconstruct a near complete global phylogeny of Viviparidae. Molecular dating and ancestral range estimation were undertaken to obtain the time frame for the dispersal events. Results from these analyses were contrasted with paleoclimate and paleogeology to better understand the biogeography of Indian viviparids. Results support at least two dispersal events into India from Southeast Asia. The earlier event is likely to have occurred during a warm and humid Eocene period before a permanent land connection was established between the two landmasses. While the more recent dispersal occurred post-suturing and overlapped with a time in late Tertiary to Quaternary when arid climate prevailed. However, we could not firmly establish how the marine transgressions influenced the dispersal events. Even though most biotic exchange between India and SEA are noted to be post-suturing, our results add to a growing body of work that suggests faunal exchange pre-suturing probably mediated by intermittent land connections.

## 1. Introduction

Historical biogeographic patterns are governed by factors such as continental movements, climatic fluctuations, and, the ecology of organisms (Brown and Lomolino, 1998). Biogeography of the Indian subcontinent, which falls under the Indian subregion (IS) (Wallace, 1876), offers us a unique opportunity to understand these factors given the rich history of geological and climatic events in the subcontinent (Valdiya, 2010). In recent times many authors have stressed the importance of the ‘into-India’ scenario where flora and fauna colonized IS from Asia. Molecular data has also supported this scenario in a suite of organisms of different taxonomic groups (Agarwal et al., 2014; Datta-Roy et al., 2012; Klaus et al., 2016 and the references therein). However, the relative importance of various paleoclimatic and paleogeological factors with respect to the ecology of the species is not well understood.

The dispersal rates into IS from various adjoining landmasses varied across time due to fluctuations in climate and geographical connectivity (Klaus et al., 2016), although the influx began from about 70–65 mya (Valdiya, 2010). One reason behind this is the complicated history of India-Asia collision for which there are several competing theories. According to one, Indian plate collided with the Asian plate at around 50 mya (Royden et al., 2008). The collision led to the formation of a depression called the Himalayan foreland basin which underwent a marine transgression soon after, lasting till 31 mya (Bera et al., 2008; Singh et al., 2016; Valdiya, 2010). According to another theory, Indian plate glanced across parts of Southeast Asia (SEA) around 55 mya and the final suturing began at around 35 mya. All this while Neotethys Ocean lay between the two landmasses and they were only connected intermittently (Aitchison et al., 2008). In both cases, the presence of the sea might have acted as a barrier to dispersal for saltwater intolerant taxa up until 35–31 mya.

The Bengal basin forms a large part of the dispersal corridor between Southeast Asia (SEA) and IS. Geological evidence suggests that recurring marine transgression events have taken place in the Bengal basin starting from Eocene. The transgression events might have acted as a barrier to dispersal even after formation of a continuous land bridge between the two landmasses. To elaborate, there was a transgression event in the basin during Eocene, followed by a regression during Oligocene. There were repeated cycles of transgression and regression during Miocene. The sea finally retreated during the early Quaternary (Alam et al., 2003; Banerji, 1984). However, there is some evidence of marine conditions prevailing in some parts till late Pleistocene (Roy and Chatterjee, 2015).

Lastly, post-collision the IS (55 mya–present) experienced a series of climatic upheavals. In the early stages of the collision, throughout Paleocene, early and mid-Eocene warm and humid conditions prevailed in IS which facilitated the growth of warm tropical forests (Kar, 1985). The landmass subsequently turned into a much drier place as evident from vast expanses of scrubs and grasslands in a large part of peninsular India. The aridification is largely linked to intensification of the monsoon climate and the onset of seasonality. Although the earliest evidence for monsoon climate comes from early Eocene (Licht et al., 2014), the first major aridification event took place during the Eocene-Oligocene boundary, also known as Eocene Oligocene Transition (EOT), due to a global cooling event (Zachos et al., 2001). Palynological studies suggest that EOT is also concurrent with a reduction in the warm tropical forest cover and establishment of regional variation in floral elements. Later on, evidence for monsoon climate and seasonality is found ~24 mya (Clift et al., 2008). Early Miocene onwards, a shift towards warmer and humid climate is reported, which culminated in mid-Miocene climatic optimum. As a consequence, a rainforest belt was established from SEA to IS which facilitated dispersal of wet-adapted Southeast Asian groups to IS. Klaus et al., 2016 reports dispersal rate from Southeast Asia to India approaching a peak during this time (21–11 mya). Warm, humid climate is also conducive for dispersal of freshwater groups (Klaus et al., 2010). The last stages of aridification began during the late Miocene (Dettman et al., 2001; Molnar et al., 1993; Molnar & Rajagopalan, 2012; Nelson, 2007). The late Miocene aridification resulted in the expansion of C4 grasslands and diversification of several arid-adapted species groups (Agarwal & Ramakrishnan, 2017; Deepak & Karanth, 2017). This event, in particular, is also suggested to have decreased the rate of dispersal of wet adapted species from Southeast Asia (Klaus et al., 2016).

Aridification leads to paucity of waterways and available routes for colonization of a new place. This restricts the dispersal or range expansion of freshwater organisms (Daniels et al., 2006; Unmack et al., 2012). A marine strait can also act as barrier owing the inability of freshwater organisms to withstand salinity. Freshwater gastropods are excellent model systems to understand such patterns owing to their limited dispersal ability. Till date, only one study has shown the ‘into-India’ scenario in a freshwater snail (Köhler and Glaubrecht, 2007), however, this study was based on limited sampling from India and lacked molecular dating. Another interesting aspect of Indian biogeography is the endemic radiations resulting from the insular nature of the subregion (Karanth, 2015) However, the degree to which organisms showcase such patterns is contingent on the factors governing the dispersal events into and out of the IS. Since India has around 13 freshwater snail families (Rao, 1989), much of the biogeographic history remains unexplored. Hence, a study similar to this has the potential to provide new insight into the evolution of biodiversity in the subcontinent in the context of paleogeology and paleoclimate.

The freshwater snail family Viviparidae Gray, 1847 is present on every continent except Antarctica and South America. Out of the three subfamilies that belong to Viviparidae, two are found in North America and Europe; part of the supercontinent Lauria. The only subfamily present in IS, Bellamyinae, is also distributed in parts of East and Southeast Asia, Australia and Africa. Much of the generic diversity of this subfamily is from East and Southeast Asia. A study in 2009 targeting largely African taxa, showed *Bellamya bengalensis*, a species described from IS and parts of Southeast Asia, nested within a larger Southeast Asian group suggesting colonization of IS by Viviparids from SEA (Sengupta et al., 2009). However, in absence of thorough sampling and molecular dating, the exact number of dispersals and factors governing those events remains uncertain.

In this study, we combine robust phylogenetic, molecular dating and biogeographic analysis to address four important questions: 1) Are the *Bellamya* species distributed in IS part of an endemic IS radiation or have there been multiple colonizations? 2) Whether Viviparidae dispersed into IS before or after suturing of Indian plate (approximately 34–31 mya) with SEA. 3) Did the dispersal event(s) predate, postdate or overlap with any of these paleoclimatic events: EOT (~34 mya), mid-Miocene climatic optimum (18–15 mya) and late Miocene intense aridification (~10 mya onwards). 4) Did the dispersal event(s) occur during one of the many marine regression events in the Bengal basin?

## 2. Materials and Methods

### 2.1 Taxon Sampling

Whole animal samples were collected from outside protected areas which do not require acquiring a permit from the authorities and preserved in absolute ethanol. All the described species belonging to family Viviparidae from IS were sampled from or near their type locality. Three species of Viviparid snails are distributed in the IS: *Bellamya bengalensis*, *Bellamya dissimilis* and *Bellamya crassa*, however, their taxonomy is dubious. Since, *B. crassa* is sometimes considered as a junior synonym of *B. dissimilis* (Rao, 1989) and there were not many diagnostic characters to distinguish between *B. dissimilis* and *B. crassa*, we referred to those morphotypes as *B*. cf. *dissimilis*. Northeast India (NEI) are considered to be part of Indo-Chinese subregion, which includes parts of SEA as well (Barley et al., 2015; Elwes, 1973; Mani, 1974; Wallace, 1876). Since, both the species are known from SEA as well, we carried out sample collection from NEI and considered those individuals as representatives of *B. bengalensis* and *B*. cf. *dissimilis* from SEA. We have also collected samples of other genera that belong to Viviparidae from NEI (see Table A1 for a complete list of sampled individuals and Figure A1 for a map of sampling locations in Appendix A). Sequences of most of the Bellamyinae genera distributed in Asia and Africa and a few species belonging to subfamilies Viviparinae and Lioplacinae were generated by Sengupta et al. (2009) were obtained from GenBank (see Table A2 in Appendix A). Two species from sister family Ampullariidae served as outgroup following previous studies (Sengupta et al., 2009).

### 2.2 Gene Sampling

DNA was extracted using Qiagen blood and tissue extraction kit from foot muscle tissue of gastropod samples. Extracts were quantified using nanodrop, PCR amplified, purified and sequenced. A total of three nuclear and two mitochondrial genes, consisting of 1065 and 896 number of nucleotides respectively, were sequenced (see Table 1).

**Table 1:** A list of genes used in the study, their abbreviations, the primers used and the references.

| **Gene Name** | **Abbreviation** | **Primers** | **Reference** |
| --- | --- | --- | --- |
| 16S ribosomal RNA | 16S rRNA | 16Sar-L and 16Sbr-H | (Palumbi, S.R., Martin, A., Romano, S., Mcmillan, W.O., Stice, L., Grabowski, 1991) |
| Cytochrome c oxidase subunit I | COI | COIF ASMIT1 and External b | (Schulmeister et al., 2002; Stothard and Rollinson, 1997) |
| 28S ribosomal RNA | 28S rRNA | C1 and R2 | (Mollaret et al., 1997; Morgan et al., 2002) |
| 18S ribosomal RNA | 18S rRNA | 18SYLMFOR and 18SYLMREV | (Stothard et al., 2000) |
| Histone H3 | Histone H3 | H3F and H3R | (Colgan et al., 1998) |

### 2.3 Phylogenetic Analysis

The downloaded and amplified sequences are aligned using MUSCLE in Mega 7 (Kumar et al., 2016). Initially, the mitochondrial and nuclear genes were analyzed separately and thereafter concatenated for further analysis. The protein-coding genes were translated into amino acid sequences to investigate the probable presence of pseudogenes. The best partition scheme and models of sequence evolution were determined using Bayesian Information Criteria in PartitionFinder2 (Lanfear et al., 2017). Partitioning scheme and models of sequence evolution were assessed twice, once for phylogenetic analyses and again for molecular dating owing to the difference in models of sequence evolution allowed in different softwares used (see Table A3 in Supporting Information for a full list of partitions and models used). Maximum likelihood (ML) analysis was carried out using RAxML HPC 8.1.2 (Stamatakis, 2014) implemented in raxmlGUI 1.5 (Silvestro and Michalak, 2011). Ten ML searches were run along with thorough bootstraps with 10,000 replications. We followed the partition scheme suggested by the PartitionFinder analysis. The Bayesian analysis was implemented in MrBayes 3.2 (Ronquist and Huelsenbeck, 2003). Two independent MCMC chains were run for 5000000 generations and sampled every 500 generations. The lowering of the standard deviation of split frequency to below 0.01 was used as a means for determining convergence. Additionally, we checked whether all parameters have reached stationary phase and whether the sampling was thorough (>200 ess values) in Tracer 1.5 (Rambaut, 2009). The first 25% of the samples were discarded as burnin.

### 2.4 Species delimitation

The two *Bellamya* species exhibited high intraspecific genetic variation in the mitochondrial DNA, suggesting that they might be species complexes with many cryptic species. To ascertain the number of dispersals into India, the actual number of species per Bellamya lineage from IS and their distribution must be known. Therefore, we carried out independent species delimitation analysis on the two species groups *Bellamya bengalensis* and *B.* cf*. dissimilis*. We adopted two single locus-based approaches: Poisson Tree Process (PTP) and multi-rate Poisson Tree Process (mPTP). PTP utilizes the phylogenetic species concept and incorporates the number of substitutions directly to infer the number of putative species (Zhang et al., 2013). mPTP is an improvement over PTP in that it incorporates different levels of intraspecific genetic variation (Kapli et al., 2017).Bayesian trees generated through MrBayes were used as inputs. Four independent chains were run for 100000000 generations for both the PTP and mPTP analyses and sampled every 10,000 generation. The convergence of the chains was confirmed by the decrease in average standard deviation of delimitation support values (ASDDSV). First 25% of the samples were discarded as burnin.

### 2.5 Molecular Dating

A time tree was reconstructed, from the concatenated dataset with all the sampled individuals using BEASTv1.8.3 (Drummond et al., 2012). The partitioned dataset was assigned separate models of sequence evolution following the suggestion from the PartitionFinder analysis and unlinked relaxed lognormal clock models. A yule speciation prior was used as the tree prior. Initially, we used an external molecular clock (Wilke et al., 2009). However, the age estimates of the older nodes were much younger than as suggested by the existing fossil record. Hence, two fossils calibration were employed to estimate the divergence dates. 1) The oldest fossil recognized as *B. bengalensis* was used to calibrate the stem *B. bengalensis* lineage (Lognormal prior, Offset = 2.3 mya, Log (Mean) = 0.5, Log(Stdev) = 0.5). 2) The stem African lineage was calibrated with the oldest African fossil (Lognormal prior, Offset = 19 mya, Log (Mean) = 1.36, Log(Stdev) = 0.63) (see Appendix B for a detailed discussion on usage of these calibrations). A broad CTMC ref rate reference prior (Minin and Suchard, 2007) was set on all the partitions. Two independent MCMC runs were performed for 100 million generations with sampling every 10,000 generations and later viewed in Tracer 1.5 (Rambaut, 2009) to assess convergence of the runs. The first 25% of the samples were discarded as burnin and the rest were summarized in TreeAnnotator 1.8.0 (Rambaut, & Drummond, 2013).

### 2.5 Ancestral Range Estimation

Ancestral Range Estimation was performed using dispersal-extinction-cladogenesis + jump dispersal (DEC+J) ML model implemented in R package BioGeoBears 1.1 (Matzke, 2013) with the intent of understanding the range evolution of the taxa belonging to subfamily Bellamyinae and time and a number of colonization events into IS. The maximum clade credibility tree, generated after summarizing the BEAST output, was edited to include only the members of subfamily Bellamyinea, using the package APE (Paradis et al., 2004) in R 3.4.2 (available at https://www.r-project.org/). The whole distribution range of Bellamyinea was divided into four regions: 1) IS, 2) SEA including NEI, Southern China, and Japan, 3) Africa, and 4) Australia. The areas roughly correspond to different biogeographic regions and sub-regions (Elwes, 1873; Mani, 1974; Wallace, 1876). Moreover, many of these areas harbor endemic radiations by virtue of which they can be called separate biogeographic unit. 1) dispersal from Australia to India and Africa and vice versa, and 2) dispersal between SEA Africa were also assigned a low probability (0.01). Bellamyinae-like fossils have been retrieved from parts of middle-east, thus, it is likely that the snails have dispersed to Africa through India and Middle East (Ashkenazi, Klass, Mienis, Spiro, & Abel, 2010; Heller, 2007). Therefore, we assigned a probability of one to dispersal between India and Africa although they are not adjoining areas. Two independent analysis with slightly different configuration were run: 1) We pruned the *B. bengalensis* and *B.* cf*. dissimilis* clades to include only one individual per putative species as inferred from PTP analysis. The distribution range of putative species from NEI and IS were coded as SEA and India respectively. 2) As mPTP analysis suggested that both the aforementioned clades constitute only one species each: we pruned the dataset to retain only one individual each from *B. bengalensis* and *B* cf*. dissimilis* clade. The distribution range was coded as Southeast Asia-India.

## 3. Results

### 3.1 Phylogenetic analysis

The ML and Bayesian trees obtained from the concatenated dataset were congruent with each other (see Figure 1). The combined dataset was similar to the mitochondrial tree, as it was driven by the high number of changes in the mitochondrial dataset. Since the nuclear tree was largely unresolved due to low variation, we used the topology based on the combined dataset for further analyses. The subfamily Bellamyinea was obtained monophyletic in accordance with previous studies (Sengupta et al., 2009). Two major clades were retrieved within Bellamyinea: A, consisting solely of Southeast Asian taxa; B, consisting of taxa from SEA, Australia, Africa, and IS. *Bellamya bengalensis* and *Bellamya* cf*. dissimilis* were not found to be sisters to each other. The African species were obtained as sister *to B.* cf. *dissimilis* clade whereas Southeast Asian genus *Filopaludina* was sister to *Bellamya bengalensis*. Both the Indian lineages were nested within a greater Southeast Asian radiation. In a previous study, wherein *B.* cf*. dissimilis* from India was not sampled, the African clade was sister to the Southeast Asian clade (Sengupta et al., 2009). The individuals sampled from Lake Malawi in the rift valley of Africa (*Bellamya jeffreysi* and *B. robertsoni*) were monophyletic as previously shown (Sengupta et al., 2009).

**Figure 1:**
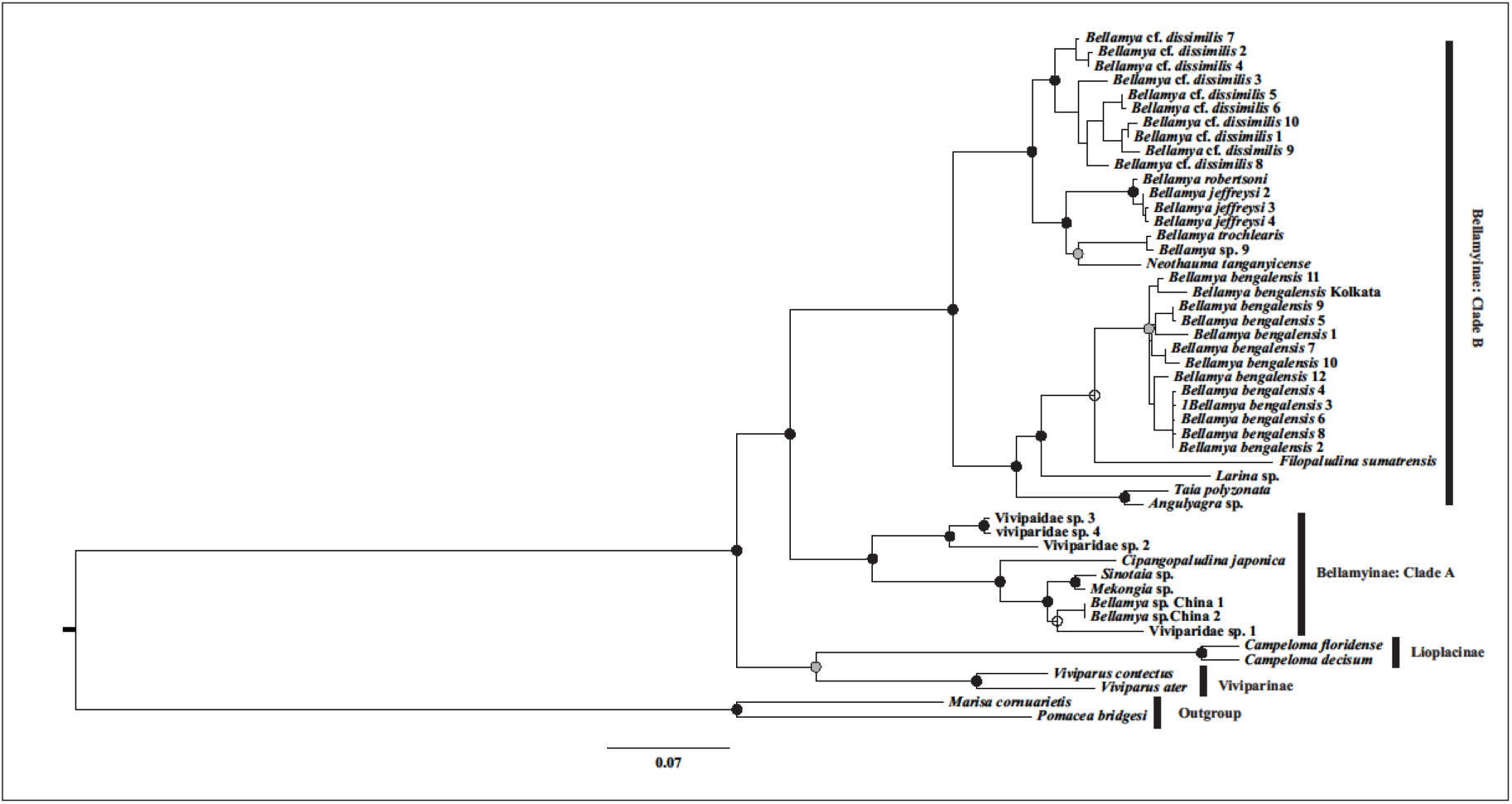
Maximum likelihood and Bayesian phylogeny of Viviparidae (concatenated data: COI, 16S, 18S, 28S, Histone H3). Nodes marked with black circles indicate high support (bootstrap value: ≥ 80; Bayesian posterior probability ≥ 0.95); nodes with open circle represent low bootstrap (< 80) but high posterior probability; nodes with gray circle indicate high bootstrap but low posterior probability (< 0.95). Multiple samples of the same species are tagged with serial numbers (eg. *Bellamya bengalensis* 1, see SI Table 1). Colors have to be used for this figure in print.

### 3.2 Species Delimitation

The PTP analysis retrieved several species in both the lineages. The results suggest that *B.* cf*. dissimilis* consists of six species while *B. bengalensis* has eight species. However, the mPTP analysis suggested that each clade consists of only one species. Previous studies have incorporated other lines of evidence such as morphology, morphometry, and environmental data in order to determine the correct number of species (Joshi & Karanth, 2012; Karanth, 2017; Lajmi et al., 2016). In the absence of additional lines of evidence, we decided to carry out independent Ancestral Range Estimation analysis, following the results of PTP and mPTP analysis respectively, to ascertain if the biogeographical inferences differ drastically between the two speciation models.

### 3.3 Molecular Dating

According to the divergence dating analysis, the two major clades in the Bellamyinea phylogeny, A and B diverged from each other on a date ranging from mid-Cretaceous to Eocene (78.8–41.8 mya 95% HPD) (see Figure 2). *Bellamya bengalensis* lineage has diverged from its sister *Filopaludina* relatively recently (13.1–5 mya). The clade consisting of *B.* cf*. dissimilis* and the African viviparids diverged quite early from their Southeast Asian sisters (50.2–27.6 mya) compared to *B. bengalensis* clade. The *B.* cf*. dissimilis* clade diverged from their sister clade, the African viviparids around 24.5–19.6 mya.

**Figure 2:**
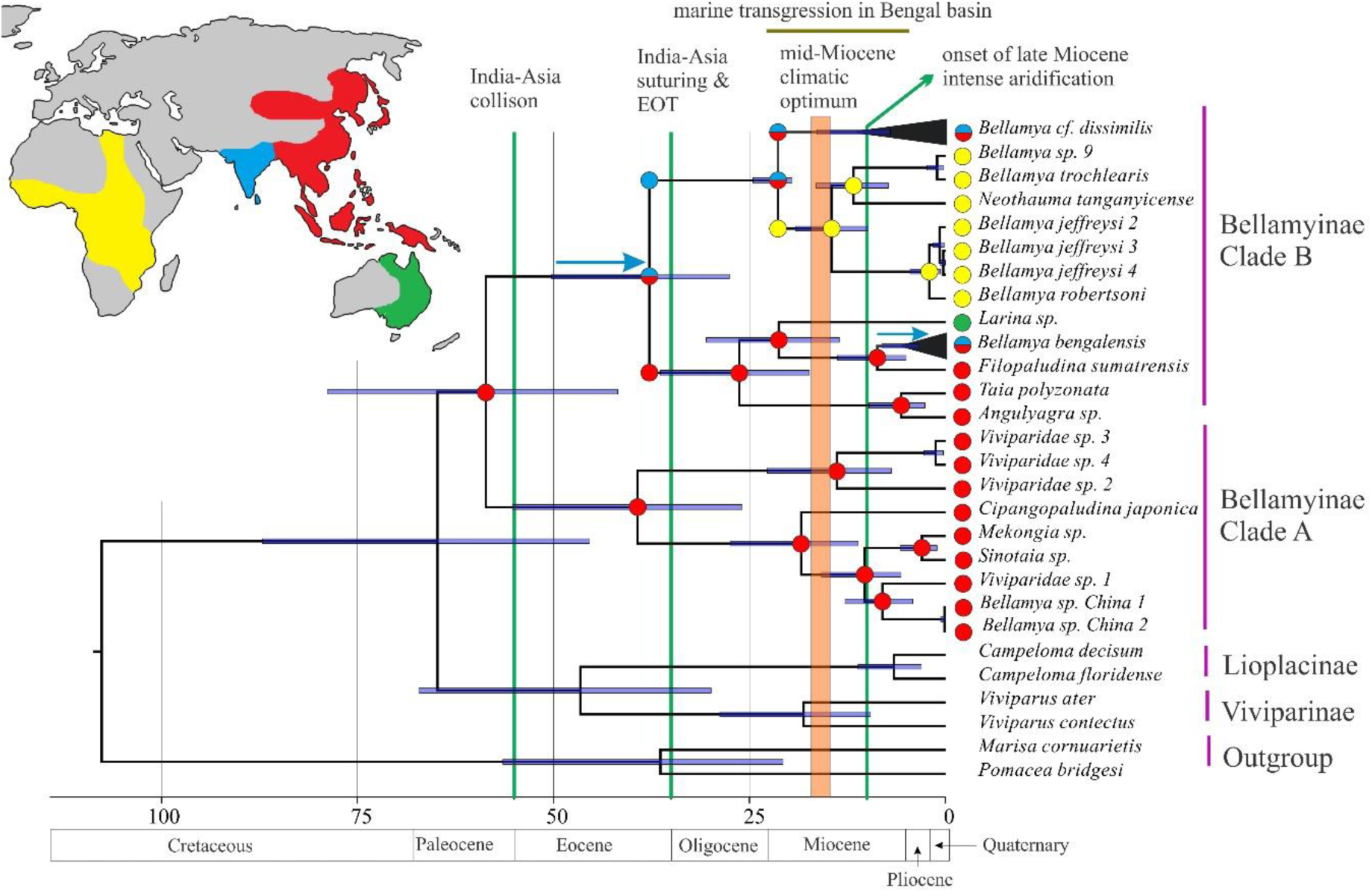
Results of divergence dating and ancestral range estimation analysis following mPTP species delimitation. The colored circles represent the ancestral areas reconstructed at respective nodes and their inheritance. The colors correspond to the color of the biogeographic areas used in the analysis as seen in the map on the left. The various vertical and horizontal lines, and the shaded area refer to paleogeological and paleoclimatic events of interest. The horizontal bars at the nodes of the phylogeny represent 95% HPD of the divergence dates at respective nodes. Colors have to be used for this figure in print.

### 3.4 Ancestral Range Estimation Analysis

The results of both the Ancestral Range Estimation analyses based on PTP and mPTP results were highly congruent with each other. Analysis based on mPTP results are shown in Figure 2. Here the most recent common ancestor (MRCA) of the ingroup (Bellamyinae) was distributed in SEA. There have been two dispersals into India from SEA, one by the *B. bengalensis* and one by the MRCA of clade B (Figure 2). The time-frame of dispersal of the latter is much older than the other and spans much of late Cretaceous to early Oligocene (78.8–27.5mya). The MRCA of *B. bengalensis* group dispersed later during late Miocene to present (13.8–0 mya). The analysis that followed the PTP results and retained multiple putative species in both the targeted lineages shows the time of the first dispersal was sometime between Early Eocene to early Oligocene (50.2–27.6). The difference in inferred dates is largely owing to the different processes behind the colonization of India inferred by these two analyses (Cladogenesis through sympatry vs. jump dispersal). *B. bengalensis* clade however colonized IS multiple times between is between (3.5–0) mya (see Figure A2 in Appendix A). A more rigorous species delimitation analysis that includes multiple lines of evidence will help resolve the conflict. However, as we discuss below, these deviations do not alter the overall inference of the analyses.

## 4. Discussion

This study aims to understand how the dispersals of Viviparid Bellamyinid snails from SEA into IS was shaped by the geological and climatic history of the region. We demonstrate that the distribution of the root node of Bellamyinae was in Southeast Asia. This is in agreement with a higher generic diversity of Bellamyinae taxa in Asia and presence of Southeast Asian species in both clade A and B. The distribution of the sister subfamilies Lioplacinae and Viviparinae in Europe and North America respectively and overall Laurasian distribution of the parent family also supports a Southeast Asian rather than Indian or African origin. The study further showed that there were two independent dispersal events from SEA into IS at varying time points (from 78.8–27.5 mya and 13.8–0 mya respectively). These time frames overlap with many paleogeological and paleoclimatic events of importance, and the role of these in governing the dispersal events will be discussed in the following sections. The two independent dispersal events giving rise to the two described Bellamyinid species distributed in IS also showed that they are not part of an endemic IS radiation.

### 4.1 ‘Into-India’ dispersal of Viviparidae

Previous studies suggested that formation of a permanent subaerially exposed land connection between Indian and Asian plate have resulted in marked increase in the dispersal rates between India to Southeast Asia after 40 mya (Klaus et al., 2016). Particularly, limnic organisms are least expected to cross the oceanic barriers between the two landmasses before the establishment of a land connection. However, there are instances where freshwater organisms managed to cross over even before the beginning of suturing of the two plates i.e. formation of a permanent land connection (Klaus et al., 2010, Li et al., 2011). This study suggests that two dispersal events into IS from SEA have taken place, both of which are inferred in clade B. The younger dispersal event took place post-suturing of the two landmasses i.e. after 34 mya. However, the time window inferred for the older dispersal (from 78.8–27.5 to 50.2–27.6 mya) span from late Cretaceous to mid-Oligocene (See Figure 2 and Figure A2 in Appendix A). Although a post-suturing dispersal scenario also cannot be rejected, much of this time window spans the pre-suturing time period, thus a pre-suturing dispersal scenario could be more likely. Given that long-distance transoceanic dispersal is unlikely for limnic organisms, how do we explain the possible pre-suturing dispersal? According to Acton’s model, the drifting Indian plate made its initial contact with the Asian plate at 55 mya and there might have been intermittent land connections till the suturing between the two plates began (~34 mya) (Aitchison et al, 2008). Thus, although other possibilities cannot be ruled out until more precise divergence dates are obtained, an Eocene dispersal for the MRCA of clade B seems to be a more likely scenario. Other studies have also reported similar results in a variety of taxa (Kluas et al., 2010; Li et al., 2013).

This brings to fore the role of past climate change on the dispersal of Southeast Asian gastropods into India. As we have pointed out in the previous paragraph, much of the dispersal window for the MRCA of clade B is pre-suturing. This was also the time when much of IS and SEA was covered with megathermal wet forests (Morley, 2000; Morley, 2003). Previous studies have also suggested that Eocene wet forests have facilitated India-SEA dispersal of floral and freshwater elements (Klaus et al., 2010; Morley, 2000; Morley, 2003). Furthermore, EOT that took place around 34 mya is known to have led to range fragmentation and widespread extinction in wet adapted forest-dwelling species (Gower et al.,2016; Agarwal et al., 2015). The aridification was also likely to lead to loss of a continuous freshwater dispersal corridor, thus making dispersal post-EOT less possible. Although the lineage persisted in India at least from Early Oligocene, speciation in the India clade is observed much later (16 mya onwards). This stasis also points towards either less speciation owing to lack of ecological opportunity or widespread extinction. On the other hand, the *B. bengalensis* group colonized IS from SEA after the mid-Miocene climatic optimum. Furthermore, much of the inferred dispersal window (13.8–3.5 mya or 13.8–0 mya) lie after the onset of the late Miocene aridification event (see Figure 2 and Figure S3). This suggests that at least one dispersal event was possible in spite of the aridification event. The three back dispersal events from IS to SEA, interpreted from the Ancestral Range Estimation analysis, have taken place during Pleistocene. How do we explain such exchange of freshwater fauna during this period of aridification? In this regard, it must be noted that even though there was a global shift towards colder and more arid climate, since late Miocene there have been phases of multiple reversals towards moist and warmer climates during Pliocene and Pleistocene (Kotlia et al., 1997; Sniderman et al., 2016; Wang et al., 1999). Especially during Pliocene humid and warmer phases were reestablished in many places in Asia (Passey et al., 2009; Sniderman et al., 2016; Wu et al., 2007). As seen during the mid-Miocene climatic optimum, shifts to humid climate favor proliferation of wet forests and freshwater habitats. This, in turn, could have facilitated dispersal from SEA to IS. Additionally, while many freshwater taxa depend on water flow as their most important means of dispersal, freshwater snails are known to disperse by attaching themselves to waterfowls (Kappes and Haase, 2012; Van Leeuwen et al., 2013). This particular mode of dispersal might have facilitated their dispersal even in the absence of freshwater dispersal corridors.

Another possible dispersal barrier that has seldom been factored in previously is the repeated marine transgression events in the Bengal basin. Marine transgressions are important factors creating biogeographic barriers (Bossuyt, 2014; Riddle et al., 2000 but see R. M. Brown et al., 2013). In our study, the dispersal time interval of the MRCA of clade B is such that the dispersal could have taken place during either Eocene transgression event or during Oligocene regression. Although there was a marine transgression during Eocene, there is ample evidence of dispersal in the IS-SEA boundary during that time. Thus, it is not clear how the transgression event affected the dispersal of this lineage. The colonization of the ancestors of *B. bengalensis* group, which is much more recent compared to the former, might have occurred during late Tertiary when the sea finally retreated from Bengal basin enabling them to utilize a broad terrestrial connection to disperse. Ours is the first study that addressed the plausible biogeographic implications of the aforementioned marine transgression events. However, the precise time frame of the repeated transgression-regression cycles is not known, thus limiting our understanding of the role of the marine regression event as a biogeographic barrier. Studies using multiple time-calibrated phylogenies that focus on dispersals across Bengal basin could shed some light on this.

### 4.2 Colonization of Africa

The African lineage of Viviparid appears to have colonized the continent from India. The colonization of Africa likely took place between 24.5–19.6 mya. This estimate is slightly older than the previously hypothesized time of dispersal (15-5 mya) when environmental conditions were favorable (Damme and Bocxlaer, 2009; Sengupta et al., 2009). Nevertheless, this date agrees with the oldest Bellamya-like fossil found in Africa (Pickford, 2004; Schultheiß et al., 2014).

### 4.3 Implication on taxonomy

It has previously been shown that the African, Indian and Asian members of the genus *Bellamya* are not monophyletic (Sengupta et al., 2009) In the WORMS website (available at http://www.marinespecies.org/) *Bellamya bengalensis* and *B.* cf*. dissimilis* are registered as *Filopaludina bengalensis* and *Idiopoma dissimilis* respectively. In the current study *B. bengalensis* was retrieved as sister to *F. sumatraensis*, thus supporting the change in nomenclature. However, *B.* cf*. dissimilis* has been obtained as sister to the African clade. No molecular study has been undertaken to determine the relationship between members of genus *Idiopoma* and *B.* cf*. dissimilis* to this date. This calls for a thorough and extensive integrative taxonomic review of the whole family using different diagnostic tools such as molecular evidence, morphometry, anatomy, cytogenetics and niche preference.

## 5. Conclusion

The results suggest an East and Southeast Asian origin of Bellamyinae and two independent dispersals of the described Bellamyinae species distributed in the IS. The first dispersal is likely to have occurred when Indian and Asian plates were only in intermittent contact and a warm humid climate seems to have facilitated the dispersal. However, we cannot rule out a later dispersal when the two plates were permanently sutured, and the climate changed to being cold and dry. The second dispersal occurred much after the two plates sutured and most likely during a period of dry arid climate. The role of the marine transgression in the Bengal basin in governing these dispersals remain unclear. Only recently biogeographers have begun to focus more intently on the factors governing the time of the two-way traffic between the two landmasses. Further studies addressing similar questions about Indian flora and fauna with different ecological requirements will help us better understand the relative importance of geographic and climatic factors on the evolution of Indian biota.

## Appendix A: Additional tables and figures from methods and results

**Figure A1:**
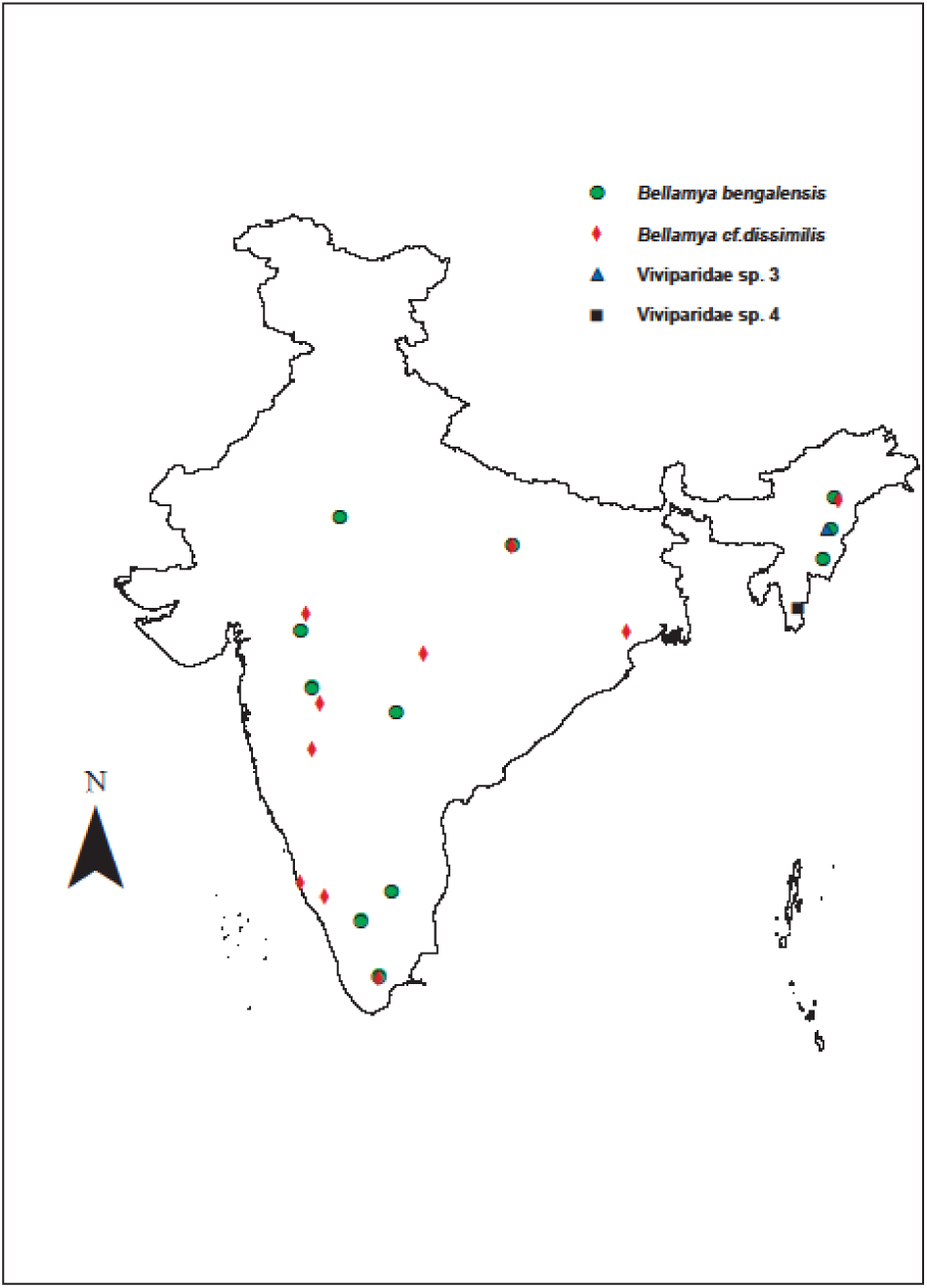
Map showing sampling locations f individual collected and amplified during the course of the study. Colors have to be used for this figure in print.

**Figure A2:**
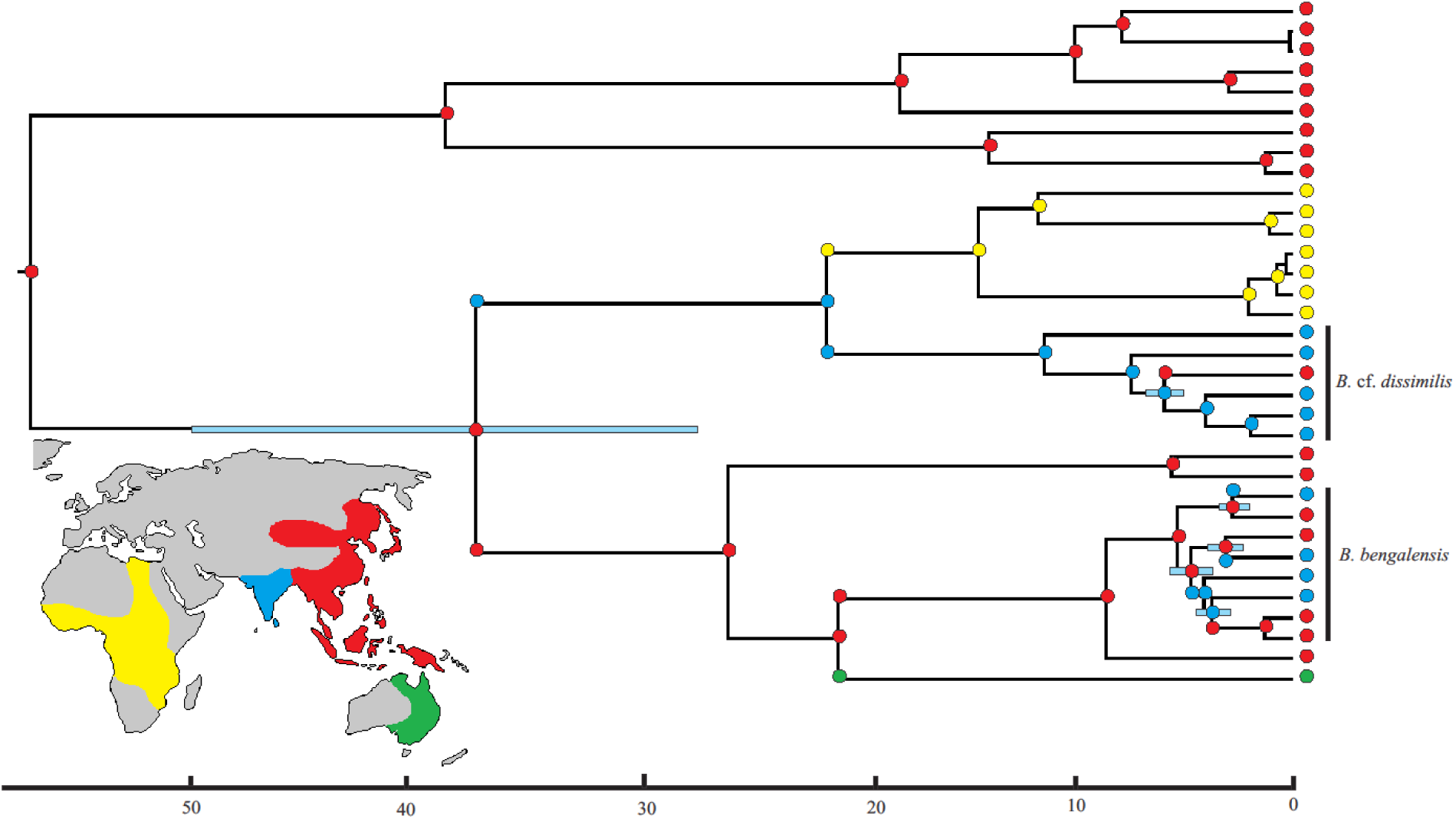
Ancestral range estimation using DEC model following PTP analysis. Discrete areas are color coded as shown in the map. Most probable range inheritance scenarios are mapped on the tree as colored circles. Tips represent present distribution of extant taxa included in the analysis. Multiple colored areas stand for distribution across different subdivisions. Colors have to be used for this figure in print.

**Table A1:**
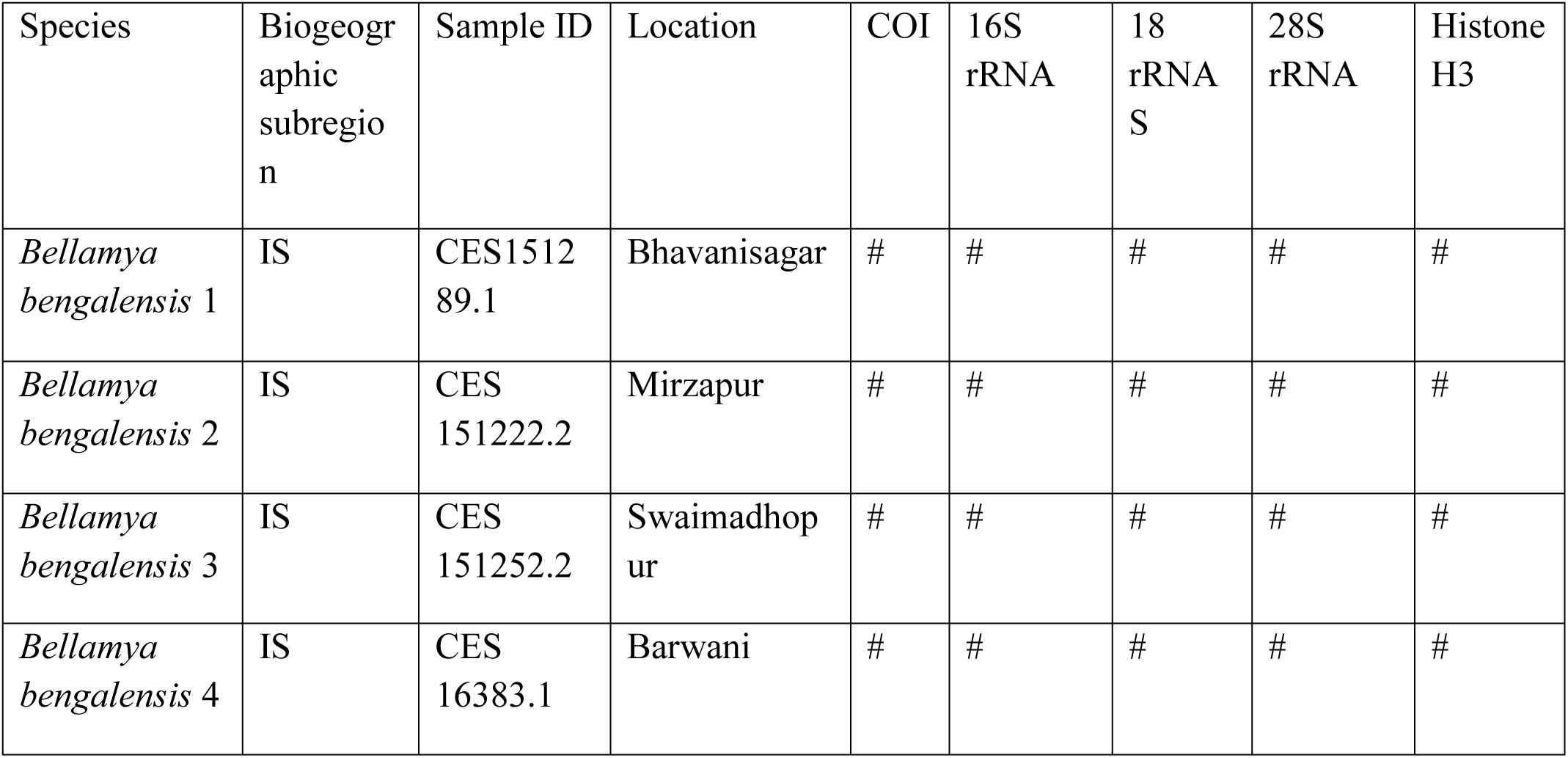
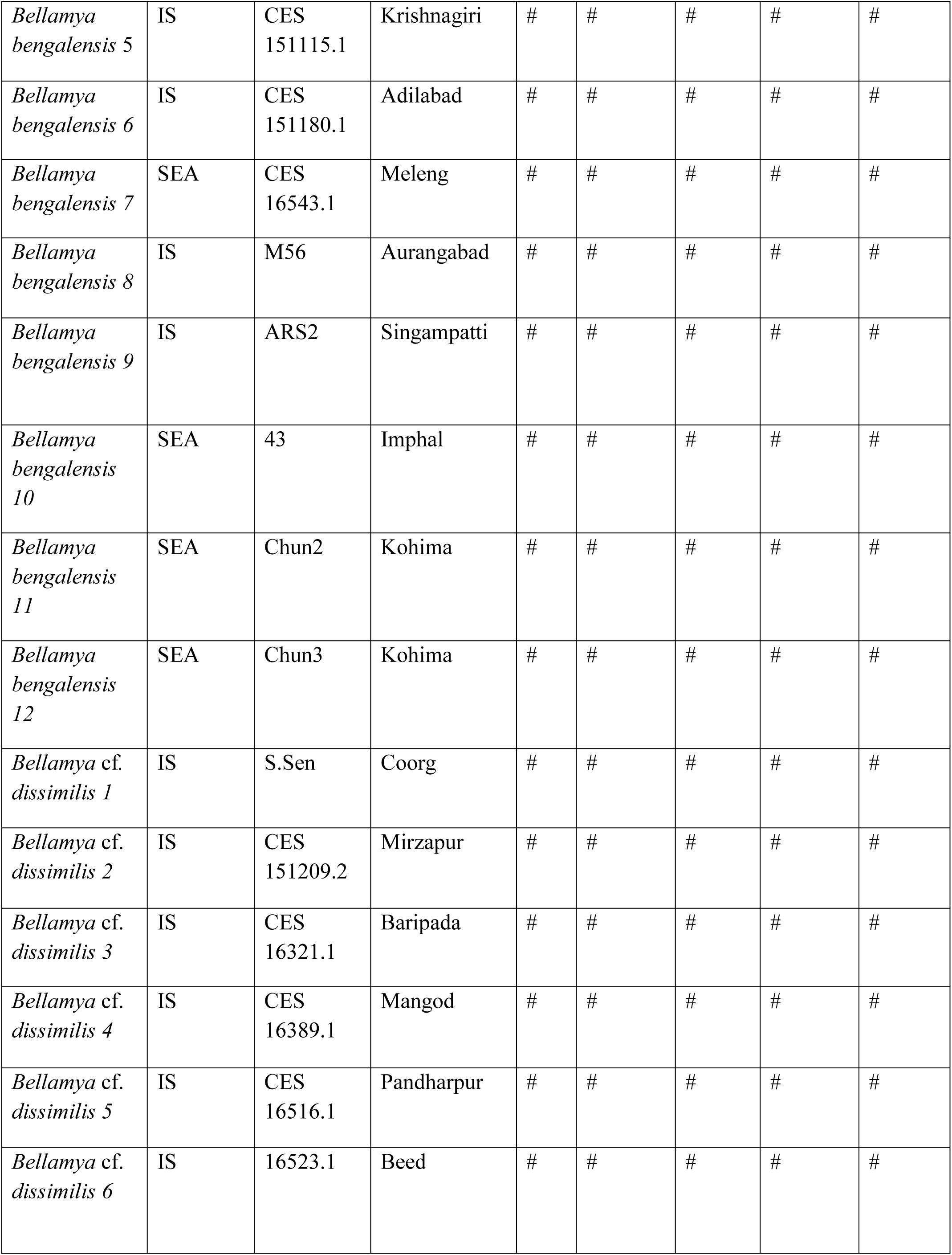
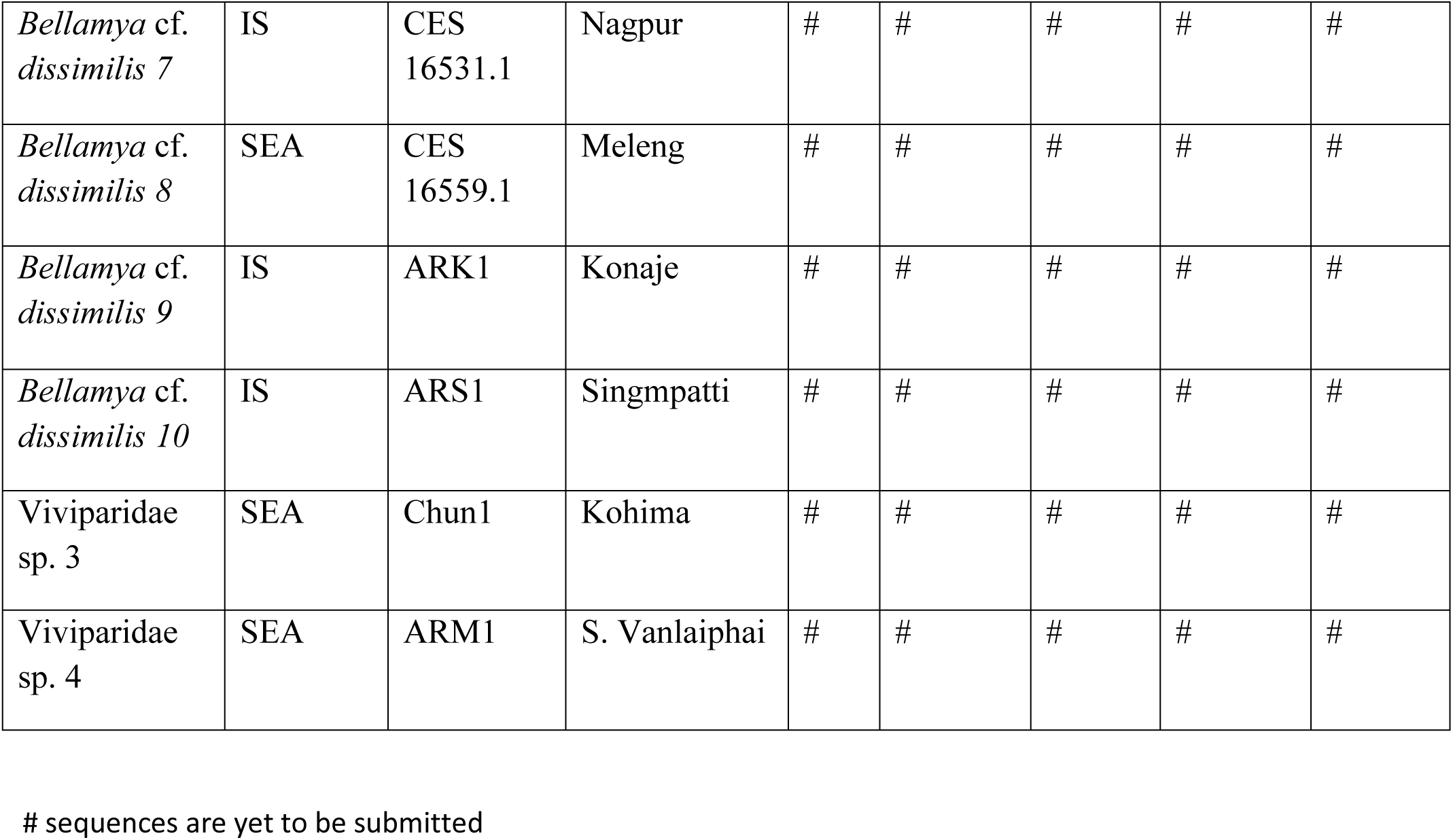
A list of specimens from India used in the study with the sampling location

**Table A2:** Sequences downloaded from genebank with accession numbers.

| Species | Country | COI | 16S | 18S | 28S | H3 |
| --- | --- | --- | --- | --- | --- | --- |
| Species distributed in SEA |  |  |  |  |  |  |
| *Angulyagra* sp | Vietnam | FJ405819 | FJ405723 | FJ405675 | FJ405628 | FJ405769 |
| *Bellamya* sp. | China | FJ405824 | FJ405725 | - | - |  |
| *Bellamya* sp. | China | FJ405826 | FJ405726 | - | - | FJ405776 |
| *Cipangopaludina japonica* | USA* | FJ405821 | FJ405736 | FJ405639 | FJ405621 | FJ405777 |
| *Filopaludina sumatrensis* | Singapore | FJ405834 | - | - | FJ405633 | FJ405770 |
| *Larina* sp. | Australia | FJ405829 | FJ405733 | FJ405681 | FJ405626 | FJ405764 |
| *Mekongia* sp. | Vietnam | FJ405820 | FJ405722 | FJ405676 |  | FJ405767 |
| *Sinotaia* sp. | Vietnam | FJ405832 | FJ405728 | FJ405688 | FJ405631 | FJ405768 |
| *Taia polyzonata* | Singapore | FJ405833 | FJ405727 | FJ405679 | FJ405632 | FJ405771 |
| Viviparidae<br>sp. 1 | China | FJ405831 | FJ405729 | FJ405687 | FJ405629 | - |
| Viviparidae<br>sp. 2 | China | FJ405825 | FJ405730 | FJ405686 | - | FJ405775 |
| Species distributed in Africa |  |  |  |  |  |  |
| *Bellamya* sp. 9 | Uganda | FJ405790 | FJ405703 | FJ405652 | FJ405611 | FJ405740 |
| *Bellamya jeffreysi* 2 | Tanzania | FJ405804 | FJ405702 | FJ405656 | FJ405583 | FJ405761 |
| *Bellamya jeffreysi* 3 | Tanzania | FJ405812 | FJ405700 | FJ405654 | FJ405584 | FJ405762 |
| *Bellamya jeffreysi* 4 | Tanzania | FJ405806 | FJ405701 | FJ405651 | FJ405585 | FJ405763 |
| *Bellamya robertsoni* | Malawi | FJ405802 | - | FJ405638 | FJ405581 | FJ405747 |
| *Bellamya trochlearis* | Uganda | FJ405816 | FJ405691 | FJ405637 | FJ405603 | FJ405741 |
| *Neothauma tanganyicense* | Zambia | FJ405788 | FJ405709 | FJ405643 | FJ405598 | FJ405739 |
| Species distributed in holarctic region |  |  |  |  |  |  |
| *Viviparus ater* | Switzerland | FJ405830 | FJ405734 | FJ405683 | FJ405630 | FJ405774 |
| *Viviparus contectus* | Denmark | FJ405778 | FJ405692 | FJ405635 | FJ405634 | FJ405737 |
| *Campeloma floridens* | USA | FJ405827 | FJ405731 | FJ405684 | FJ405624 | FJ405772 |
| *Campeloma decisum* | USA | FJ405828 | FJ405732 | FJ405680 | FJ405625 | FJ405773 |
| Outgroup |  |  |  |  |  |  |
| *Marisa cornuarietis* | South America | FJ439657 | EU274493 | EU274545 | EU274527 | EU274511 |
| *Pomacea bridgesii* | South America | FJ439658 | EU274500 | EU274553 | EU274532 | EU274514 |

\**Cipangopaludina japonica* is introduced to the USA.
- No information can be found

**Table A3:** Partition schemes and models of sequence of evolution used for different analyses

| RAXML |  | MrBayes |  | Beast |  |
| --- | --- | --- | --- | --- | --- |
| partitions | models | partitions | models | partitions | models |
| 18S rRNA | GTR+I+G | 18S rRNA | JC | 18S rRNA, 28S rRNA | TrNef+G |
| H3 codon position 1, H3 codon position 2, COI codon position 2 | GTR+I+G | H3 codon position 1, H3 codon position 2, COI codon position 2 | JC | H3 codon position 1, H3 codon position 2, COI codon position 2 | K80 |
| H3 codon position 3 | GTR+I+G | H3 codon position 3 | HKY+G | H3 codon position 3 | HKY+G |
| 16S rRNA | GTR+I+G | 16S rRNA | GTR+G | 16S rRNA | GTR+G |
| 28S rRNA, COI codon position 1 | GTR+I+G | 28S rRNA, COI codon position 1 | GTR+G | COI codon position 1 | TrNef+I |
| COI codon position 3 | GTR+I+G | COI codon position 3 | HKY+I+G | COI codon position 3 | HKY+I+G |

## Appendix B: Details on usage of fossil calibrations

The oldest *Bellamya bengalensis* fossil, unearthed from the late Pliocene deposits in Jammu (Kundal, 2013), was used to calibrate the stem lineage of *B. bengalensis*. There are reports of *B. bengalensis* fossils from contemporary or slightly younger deposits from elsewhere in the IS such as Chandigarh (Bhatia & Mathur, 1973), Saketi (Mathur, 1998), Narmada vally (Prashad, 1928). Thus, it can be safely assumed that this lineage existed in IS at that specific period. A lognormal calibration with an offset of 2.2 mya because the lineage could not be younger than the age of the fossil. The stem lineage was calibrated instead of the crown because the stem lineage might have existed much before the extant crown lineages started diversifying and the fossil could have belonged to any point in the stem lineage from the split from its sister group to the beginning of diversification in crown. Calibrating the stem will make sure all these uncertainties will be taken into consideration. To allow for the uncertainty in time estimate of the age of the stem lineage the prior was given a broad enough range (2.2–8 mya). The upper age is based on a previous dating attempt that the authors have undertaken using substitution rate of the COI gene.

The oldest African Bellamyinid fossil, unearthed from the Iriri member of the Napak formation and is dated to early Miocene (~19 mya) (Pickford, 2004). A previous study on African Viviparids used this fossil to calibrate the root node of the phylogeny consisting of African Bellamyinid species (Schultheib et al., 2014). Here we used the same logic as before that the African lieneage cannot be younger than 19 mya, and the fossil can belong to the stem lineage instead of the crown. Hence, we calibrated the stem African lineage with a lognormal prior and the offset was placed at 19 mya. The upper age of the distribution was 40 mya: the time when Africa collided with Eurasia (Van Yperen et al., 2005).

## Acknowledgements

RG Kamei and Stephen Mahony provided us samples from Kohima. Anirudha Datta Roy, Ishan Agarwal and Aparna Lajmi commented on the manuscript, and assisted during analysis. Kunal Arekar, Chinta Sidhharthan, Harshal Bhosale, Shirish Nakhale, Pradip Barua, Tarun Singh, Suhel Sheikh, Animesh Mondal, Harshil Patel, Lansothung Lotha and Rajesh Dhere helped us during sample collection. The project was funded by DBT-IISc partnership program (22-0303-0007-05-469) to PK. The fieldwork was partially supported by Rufford Small Grant for Nature Conservation (19805-1) to MS.

